# SARS-CoV-2 D614G Variant Exhibits Enhanced Replication *ex vivo* and Earlier Transmission *in vivo*

**DOI:** 10.1101/2020.09.28.317685

**Authors:** Yixuan J. Hou, Shiho Chiba, Peter Halfmann, Camille Ehre, Makoto Kuroda, Kenneth H Dinnon, Sarah R. Leist, Alexandra Schäfer, Noriko Nakajima, Kenta Takahashi, Rhianna E. Lee, Teresa M. Mascenik, Caitlin E. Edwards, Longping V. Tse, Richard C. Boucher, Scott H. Randell, Tadaki Suzuki, Lisa E. Gralinski, Yoshihiro Kawaoka, Ralph S. Baric

## Abstract

The D614G substitution in the S protein is most prevalent SARS-CoV-2 strain circulating globally, but its effects in viral pathogenesis and transmission remain unclear. We engineered SARS-CoV-2 variants harboring the D614G substitution with or without nanoluciferase. The D614G variant replicates more efficiency in primary human proximal airway epithelial cells and is more fit than wildtype (WT) virus in competition studies. With similar morphology to the WT virion, the D614G virus is also more sensitive to SARS-CoV-2 neutralizing antibodies. Infection of human ACE2 transgenic mice and Syrian hamsters with the WT or D614G viruses produced similar titers in respiratory tissue and pulmonary disease. However, the D614G variant exhibited significantly faster droplet transmission between hamsters than the WT virus, early after infection. Our study demonstrated the SARS-CoV2 D614G substitution enhances infectivity, replication fitness, and early transmission.

## Main text

The ongoing pandemic of Coronavirus Disease 2019 (COVID-19), caused by Severe Acute Respiratory Syndrome Coronavirus 2 (SARS-CoV-2), has resulted in an unprecedented impact on human society. Since its emergence in December 2019, SARS-CoV-2 has rapidly spread worldwide, causing > 26 million cases and >900 thousand deaths as of early September 2020. Compared to SARS-CoV (2003 to 2004) and Middle East respiratory syndrome coronavirus (MERS-CoV) infection (2012 to present), SARS-CoV-2 infection causes a broader spectrum of acute and chronic disease manifestations and exhibits greater transmissibility. Many patients develop asymptomatic or mild disease, but some SARS-CoV-2-infected individuals develop severe lower respiratory infections that can progress to an acute respiratory distress syndrome (ARDS), strokes, cardiac pathology, gastrointestinal disease, coagulopathy, and a hyperinflammatory shock syndrome (*1–3*). Ciliated cells in the respiratory epithelium and type 2 pneumocyte in the alveoli are the major targets of SARS-CoV-2 (*4*). Viral entry is mediated by the interaction between viral spike (S) glycoprotein and host receptor angiotensin-converting enzyme 2 (ACE2). To date, enormous efforts have focused on developing vaccines and therapeutic antibodies targeting the S protein using early ancestral isolates that have been replaced by novel contemporary strains (*5, 6*).

Pandemic spread of a virus in naïve populations may select for mutations that may alter pathogenesis, virulence and/or transmissibility. Despite the presence of a CoV proof-reading function (*7, 8*), recent reports identified an emergent D614G substitution in the spike glycoprotein of SARS-CoV-2 strains that is now the most prevalent form globally. Patients infected with the D614G-bearing SARS-CoV-2 are associated with higher viral loads in the upper respiratory tract, but not altered disease severity (*5, 9*). SARS-CoV2 S pseudotyped viruses encoding the D614G mutation were reported to exhibit increased infectivity in continuous cells lines and increased sensitivity to neutralization (*5, 10*). Structural analyses also revealed that the receptor binding domains (RBD) in the G614-form S protein occupy a higher percentage in the open conformation than the D614-form, implying an improved ability to bind to ACE2 receptor (*11, 12*). However, the D614G substitution has yet to be evaluated in the authentic SARS-CoV-2 infection models, and its function in viral pathogenesis and transmissibility remains unclear.

Previously, we generated a SARS-CoV-2 reverse genetics system based on the WA1 strain and developed primary human airway epithelial cells, human ACE2 transgenic mice, and golden Syrian hamsters as SARS-CoV-2 infection models (*4, 13, 14*). To address the function of the D614G substitution in SARS-CoV-2 replication and transmissibility, we generated an isogenic variant containing the D614G mutation in the S glycoprotein, along with a second variant that contained the nanoLuciferease (nLuc) gene in place of the accessory gene 7a (Fig 1A). To examine whether the D614G substitution enhanced authentic SARS-CoV-2 entry, four susceptible cell lines were infected with wildtype (WT)-nLuc and D614G-nLuc viruses at an MOI of 0.1. After a 1h incubation, cells were washed three times with PBS and cultured in medium containing SARS-CoV-2 neutralizing antibodies to prevent viral spreading. Luciferase signals representing initial entry events were measured at 8h post infection (Fig. 1B). In accord with pseudovirus studies (*5, 6*), the D614G-nLuc infection resulted in a 0.5 to 2-fold higher transgene expression as compared with WT-nLuc virus. Replication kinetics comparing WT and D614G viruses were performed utilizing multi-step growth curves in cell lines (Fig 1C). Although the D614G variant showed similar or slightly higher titers at the early time point (8h), its peak titers were significantly lower than the WT virus in Vero-E6 and A549-ACE2 cell lines but not in Vero-81 and Huh7. These data suggest that the D614G substitution may modesty enhances SARS-CoV-2 entry and replication in some immortalized cell lines.

**Figure 1.**
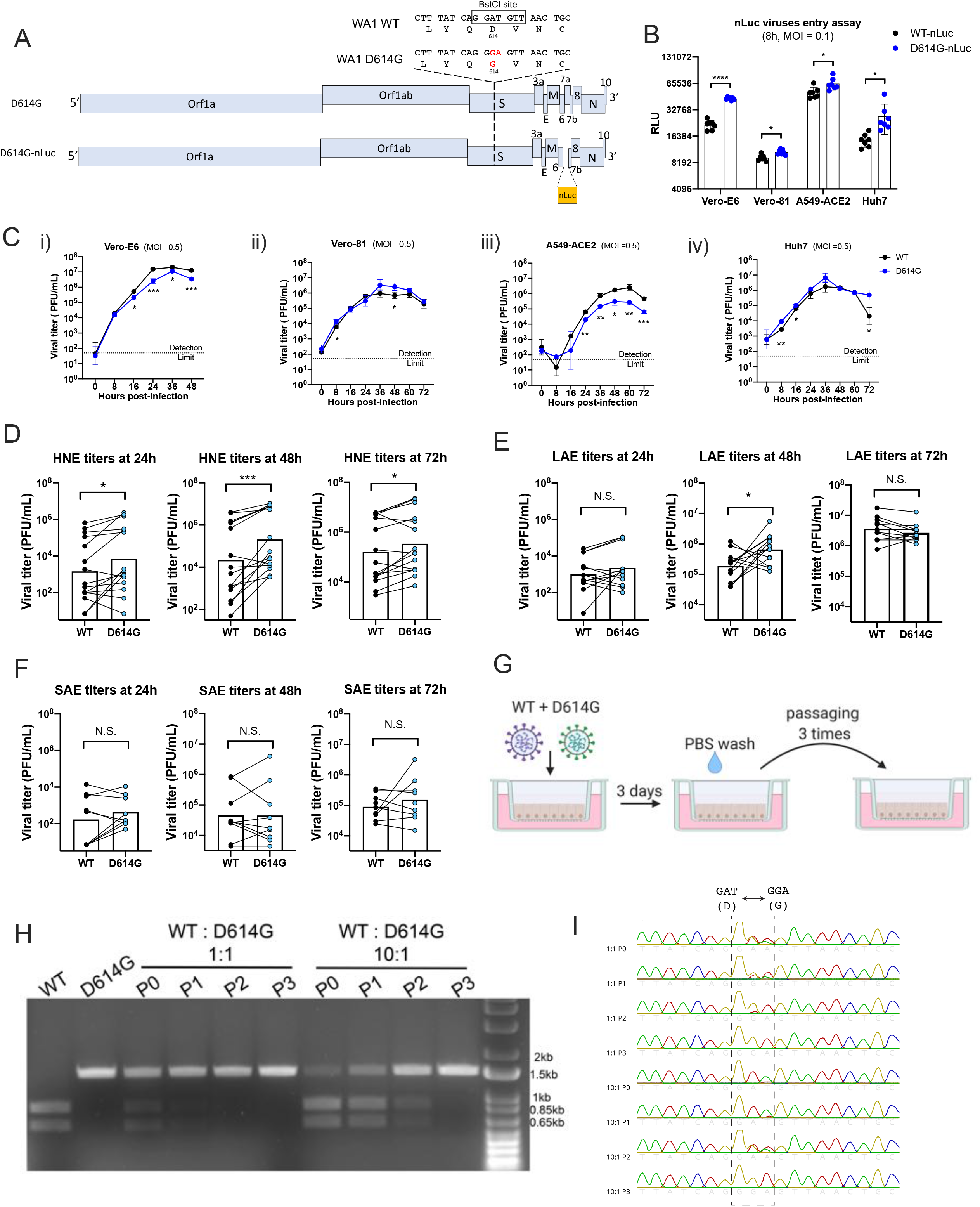
SARS-CoV-2 D614G variant demonstrate enhanced infectivity in some cell lines and replication fitness in upper respiratory epithelia. A. Genomes of recombinant SARS-CoV-2 D614G variants. B. Entry efficiency of WT-nLuc and D614G-nLuc in multiple cell lines at MOI of 0.1. After 1h infection, cells were cultured in the medium containing neutralization antibodies to minimize the secondary round of infection. The relative light unit (RLU) representing the nLuc expression level was measured at 8h post infection. C. Multi-step growth curves of the two variants at Vero-E6 (i), Vero-81 (ii) and A549-ACE2 (iii) and Huh7 (iv) cell lines at MOI = 0.5. Comparison of 24, 48 and 72h titers between the two variants infected primary nasal (D), large airway (E) and small airway (F) cells in triplicate. Triplicated titers of the two viruses in the cultures form the same donor were analyzed by paired t-test. G. Schematic of competition assay on large airway epithelial cells. Cultures were infected with 1:1 or 10:1 ratio of WT and D614G mixture at MOI at 0.5, and serially passaged three times. H. BstCI digestion of the partial S gene from the competition assay samples. A 1.5kb fragment containing the residue 614 was amplified from the total RNA collected from competition assay. I. Sanger sequencing chromatogram of S RNA collected from the competition assay. Data between the WT and D614G viruses in B and C are analyzed using unpaired t-test, and the data between the two groups in D, E and F are analyzed using paired t-test. N.S., not significantly different, *p* > 0.05; *, *p* < 0.05 **, *p* < 0.01; ***, *p* < 0.001.

Primary human airway epithelial cells from different regions of the human respiratory tract display different susceptibilities to SARS-CoV-2 infection, with the nasal epithelium being most susceptible (*4*). To evaluate the replication of SARS-CoV-2 D614G variant in the human respiratory tract, we compared WT and the D614G growth kinetics in primary human nasal epithelial (HNE) from five donors, large (proximal) airway epithelial (LAE) from four donors, and distal lung small airway epithelial (SAE) cells from three donors. Cultures from the same donor were infected with either WT or D614G virus in triplicate (Fig. 1D to E, S1A to 1B). Both viruses infect mainly ciliated cells in the primary pulmonary cultures (Fig. S1C). Paired t-test analysis suggests the D614G-infected HNE at 24, 48 and 72h, and LAE cultures at 48h exhibited significantly higher titers than cultures infected with the WT virus. This enhancement was not observed in any timepoints in distal lung SAE cultures derived from three donors. To further compare the replication fitness between the two variants in the human airway epithelia, competitive co-infection assays were performed in LAE cultures infected simultaneously with both viruses (Fig. 1G). After three continuous passages at 72h intervals, the D614G variant became dominant in the cultures regardless of whether the WT virus was at a 1:1 or 10:1 ratio over the isogenic D614G mutant (Fig. 1H and 1I). Taken together, these data suggest the D614G substitution enhances SARS-CoV-2 replication fitness in the primary epithelial cells, with a marked advantage in the upper respiratory tract epithelial cells in nasal and large (proximal) airway epithelia.

Next, scanning and transmission electron microscopy (SEM and TEM) were performed to visualize virions present on the surface of primary human airway cell cultures and did not detect significant differences in virion morphology (Fig. 2A and B). The number of spike proteins on projections of individual virions was also not significantly different between the two viruses (Fig. 2C). Further, differences in spike cleavage patterns were not observed between the two viruses in western blot analysis (Fig. 2D), in contrast to observations reported from a pseudovirus study (*15*). To compare antibody neutralization properties with reported pseudotyped virus assays (*10*), neutralization activity was measured in 10 serum samples from D614 (WT) spike-vaccinated mice using the nLuc-expressing recombinant SARS-CoV-2 encoding either WT or D614G spike. The serum samples show 0.8 to 5.1-fold higher half-maximal inhibitory dilution (ID_50_) values against the D614G virus than the WT virus, suggesting the D614G substitution rendered SARS-CoV-2 more sensitive to neutralizing antibodies (Fig. 2E and 2F). In addition, six SARS-CoV-2 RBD-binding neutralizing antibodies were evaluated and exhibited no significant difference in half-maximal inhibitory concentration (IC_50_) values against both viruses.

**Figure 2.**
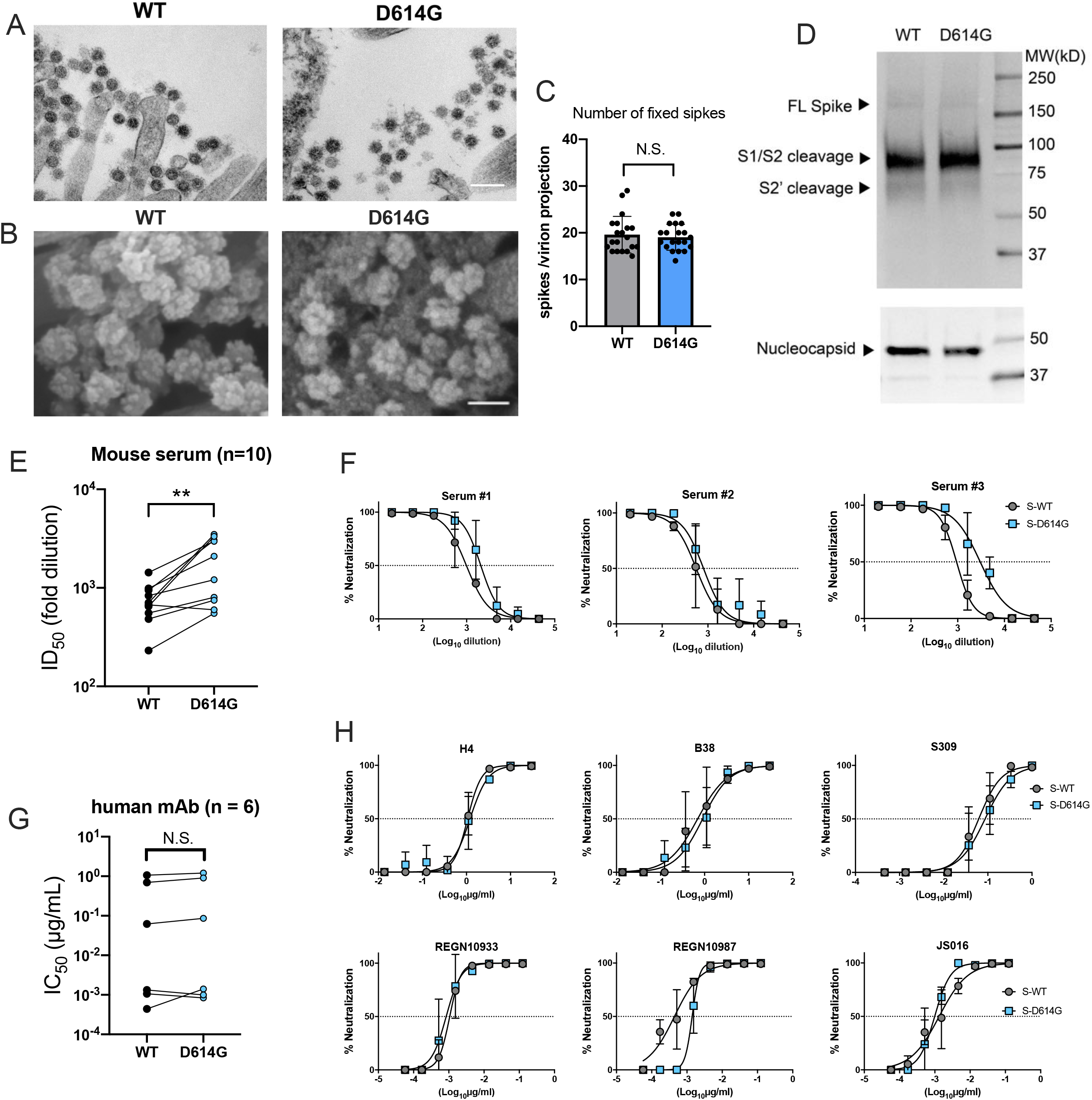
D614G substitution does not alter SARS-CoV-2 virion morphology and S protein cleavage pattern but change viral sensitivity to neutralizing antibodies. A. Transmission electron microscopy image of WT and D614G virions on airway epithelial cell surface, scale bar: 200 nm. B. Scanning electron microscopy images of WT and D614G virions on airway epithelial cell surface, scale bar: 100 nm. C. Quantification of S protein on individual SARS-CoV-2 virion projections. The number of S protein on individual virion projections from different SEM images were quantify manually, n=20. D. Western blot analysis of SARS-CoV-2 virions washed from WT-or D614G-infected LAE culture surface at 72h. Each lane contains mixed sample from triplicated cultures. Full-length (S), S1/S2 cleaved and S2’ cleaved spike protein (upper panel) and nucleocapsid protein (lower panel) were probed. E. ID_50_ values of 10 serum samples collected from D614-form Spike-vaccinated mice neutralizing WT- and D614G-nLuc viruses. F. Three representative neutralization curves of the mouse sera against both viruses. Summarized IC_50_ values (G) and individual neutralization curves (H) of 6 human nAbs against both viruses. Data between the WT and D614G viruses in E and G are analyzed using paired t-test. N.S., not significantly different, *p* > 0.05; **, *p* < 0.01.

To evaluate the function of the D614G substitution in viral pathogenesis, hACE2 transgenic mice and Syrian hamsters were infected with equal plaque-forming units (PFU) of WT or D614G viruses. Previously, we showed that SARS-CoV-2 infection in hACE2 mice exhibited a mild disease phenotype, characterized by high viral titers in the lung but minimum weight loss and undetectable nasal titers (*14*). Two groups of hACE2 mice infected with WT and D614G viruses exhibited undetectable viral titers in nasal turbinates and similar lung viral titers at day 2 and 5 post infection. One mouse (1/5) from both groups exhibited detectible viral titers in the brain (Fig. 3A). With respect to hamster studies, lung and nasal turbinate tissues collected at day 3 and 6 pi exhibited similar viral titers in each group (Fig. 3B and 3C). However, the D614G-infected hamsters lost modestly more body weight than those infected with the WT virus (Fig. 3D). Immunohistochemistry (IHC) shows similar levels of SARS-CoV-2 nucleocapsid protein in the hamster lung tissue collected at day 3, 6 and 9 from both groups (Fig. 3E, 3 G-i). Histopathological examination revealed similar severe pulmonary lesion with inflammatory cell infiltration in the alveolar walls and air spaces, pulmonary edema, and alveolar hemorrhage in both of the hamsters on day 3, extended across larger areas on day 6, and then exhibiting partial resolution by day 9 (Fig. 3F). Notably, there was no significant difference in the size of the lung lesions (Fig. 3G-ii) and the histological severity (Fig. 3G-iii). Taking together, the D614G substitution marginally enhances SARS-CoV-2 pathogenesis in the hamster, but not mouse models.

**Figure 3.**
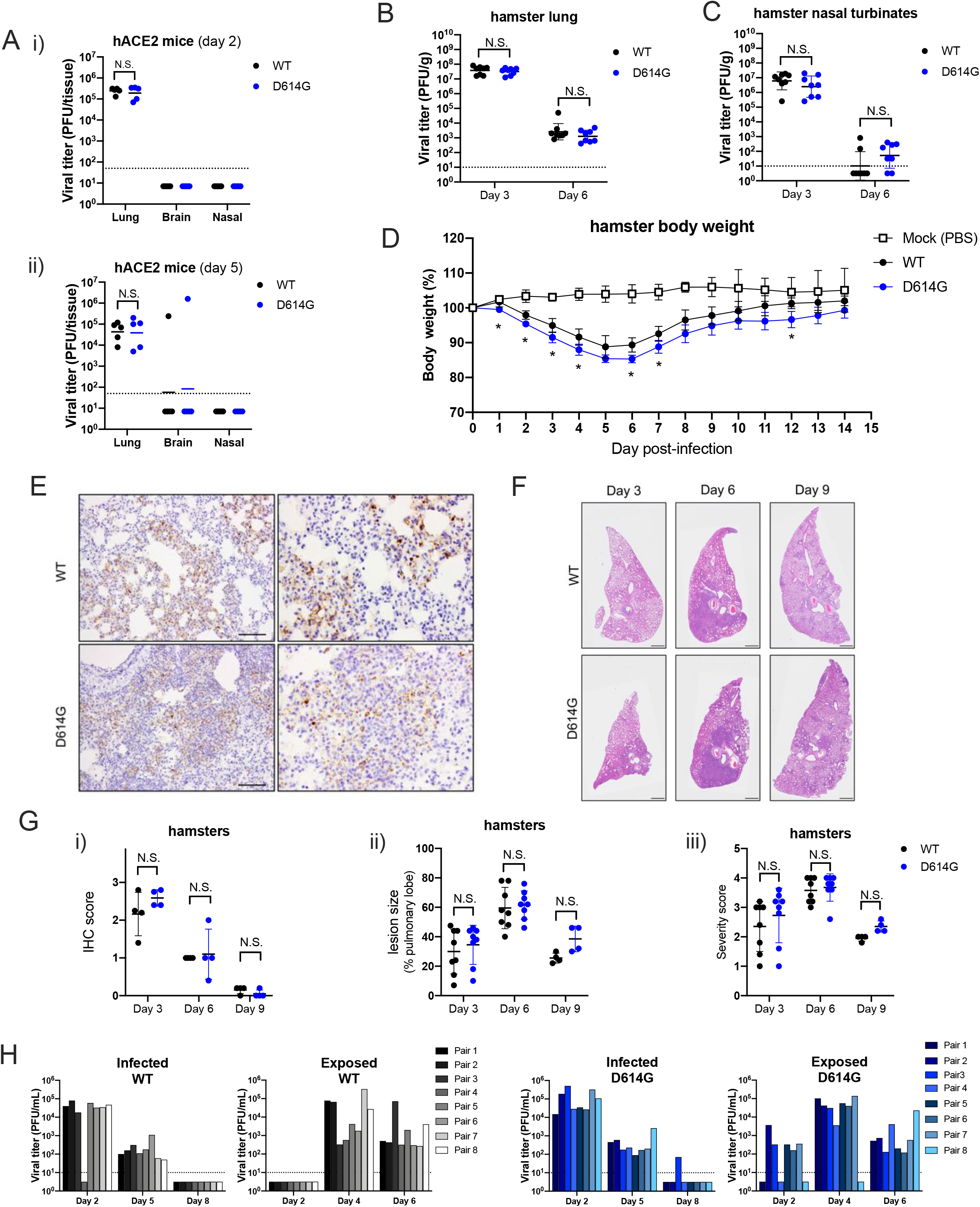
D614G variant exhibit similar pathogenesis but faster transmission than the WT virus *in vivo*. A. Lung, brain and nasal turbinate titers of WT and D614G infected hACE2 mice were determined on day 2 (i) day 5 (ii). Each mouse was infected with 10^5^ PFU of the virus, n= 5/group, plaque assay detection limit (1.7 log_10_PFU/mL) is indicated in a dash line. Viral titers of lung (B) and nasal turbinates (C) collected from SARS-CoV-2 infected hamsters at day 3 and 6. Each hamster was infected with 10^3^ PFU of virus, plaque assay detection limit (1 log_10_PFU/mL) is indicated in a dash line. D. Body weight of mock-, WT- and D614G-infected hamsters, n = 4/group. E. Immunohistochemistry (IHC) staining of SARS-CoV-2 nucleocapsid protein in the representative lung tissues collected from WT- and D614G-infected hamsters, scale bar = 100 μm. F. H&E staining of representative lung tissues collected on day 3, 6, and 9 from hamsters infected with WT or D614G, scale bar: 1mm. G-i: Quantification of IHC positive cells in hamster lung tissues, following scoring system: 0, no positive cell; 1, <10%; 2, 10-50%; 3, >50% of positive cells in each lobe of lung. G-ii. The size of pulmonary lesion was determined based on the mean percentage of affected area in each section of the collected lobes from each animal. G-iii Pathological severity scores in infected hamsters, based on the percentage of inflammation area for each section of the five lobes collected from each animal following scoring system: 0, no pathological change; 1, affected area (≤10%); 2, affected area (<50%, >10%); 3, affected area (≥50%); an additional point was added when pulmonary edema and/or alveolar hemorrhage was observed. (H) Viral titers in nasal washes collected from the infected and exposed hamster pairs in WT and D614G groups; plaque assay detection limit (1 log_10_PFU/mL) is indicated in a dash line. Data between the WT and D614G viruses in A, B, C, D, and G are analyzed using unpaired t-test. The number of transmitted hamsters at different timepoints are analyzed by Fisher exact test. N.S., not significantly different, *p* >0.05.

To evaluate the role of the D614G substitution in SARS-CoV-2 respiratory droplet transmissibility, we set up eight pairs of hamsters for each virus as described previously (*16*). Each pair comprised a naïve hamster adjacent to an infected animal 1 day after infection (Fig. S2). Viral titers in the nasal wash samples from both infected and exposed animals were measured. Both WT and D614G were transmitted efficiently to naive hamsters evident by positive nasal wash samples detected in all exposed animals at day 4 (Fig. 3H). The infected groups at all three timepoints and the exposure groups at 4 and 6 dpi exhibit similar viral titers between WT and D614G viruses. However, five of eight hamsters exposed to the D614G-infected group showed infection and detectable viral shedding at day 2 while those exposed to the WT-infected group had no infection and viral shedding (*p* = 0.0256, Fisher exact test). These data suggest the D614G variant transmits significantly faster than the WT virus through droplets and aerosols between hamsters.

Emerging viruses, like Sarbecoviruses, Alphaviruses, and Filoviruses, have undergone sequential rounds of evolution while adapting to the new human hosts in epidemic or pandemic settings (*17–19*). Among Sarbecoviruses, mutations in the Spike glycoprotein have been associated with altered pathogenesis, receptor usage, and neutralization (*20–22*), potentially challenging the development of vaccine and therapeutic antibodies that are urgently needed at present. The emergent D614G mutation in the spike glycoprotein of SARS-CoV-2 strains has raised significant concerns about potential enhancements in transmissibility, antigenicity and/or pathogenesis. Using authentic live recombinant viruses, the infectivity and fitness of D614G isogenic virus were compared in primary human cells and its pathogenesis and transmissibility were tested in hamsters and hACE2 mice. Our data unilaterally support a critical role for the D614G mutation in enhanced virus infectivity, growth and fitness in human nasal and proximal airway epithelia, but not in the lower respiratory tract airway epithelium from multiple donors. These *ex vivo* human airway culture data are consistent with the moderately increased pathogenicity, as shown by body weight changes, and improved transmission of the D614G variant in the hamster models of human disease.

Using pseudotype viruses, the D614G mutation has been suggested to increase proteolytic cleavage and S glycoprotein incorporation into virions, reduce S1 loss and promote enhanced infectivity in vitro (*5, 6, 15*). Our Western blot and SEM studies demonstrated no obvious differences in proteolytic processing or S incorporation into isogenic virions encoding the D614G mutations, perhaps reflecting differences in S trimer incorporation and presentation between authentic and pseuotyped viruses. However, our data are consistent with recent studies indicate that D614G alters spike trimer hydrogen-bond interactions, reorienting the RBD into an “up” conformation, increasing ACE2 receptor binding and infectivity (*11, 12*). Our data demonstrate that SARS-CoV2 D614G recombinant viruses are significantly more infectious in some continuous cells in culture, but more importantly, in multiple patient codes of nasal and large airway epithelial cells derived from the upper respiratory, but not lower respiratory tract. Direct competition experiments also demonstrate that the SARS-CoV2 D614G isogenic virus displays a significant advantage following passage in primary human large airway epithelial cells in vitro. Together, these data strongly support the role of the nasal epithelium and the D614G mutation in enhanced infectivity and transmission in human populations (*9*). These findings are consistent with the preferential transmission of throat SARS-CoV-2 microvariants, over sputum microvatiants in human transmission chains and in influenza virus infected ferrets (*23*).

Patients infected with the D614G variant could not be conclusively linked to increased disease severity in humans (*5, 9*). The hACE2 transgenic mouse study demonstrated equivalent virus titers in the lungs and nasal turbinates. In contrast, the isogenic D614G recombinant virus infection of hamsters resulted in significant differences in weight loss, but not pathology or virus replication in the lung and nasal turbinates. In transmission studies, the D614G isogenic was transmitted significantly faster to adjacent animals early in infection, demonstrating that the substitution preserved efficient transmission phenotypes in vivo. As SARS-CoV2 replicates preferentially in the nasal and olfactory epithelium, characterized by subtle differences driven by differences in ACE2 and TMPRSS2 cell type specific expression patterns across species (*4, 24, 25*), all of these data are consistent with a model of increased replication in the nasal epithelium and large airway epithelium, leading to enhanced virus growth and earlier transmissibility.

The effect of the D614G variant on vaccine efficacy has been of major concern. Consistent with previous studies (*10*), we demonstrated the increased sensitivity of the SARS-CoV2 D614G-nLuc variant to the antisera from D-form spike (WT) vaccinated mice. Similar findings have been reported using sera from ChAd-vaccinated mice (*26*). Together with similar neutralization properties against six SARS-CoV-2 mAbs, these data suggest that the current vaccine and mAb approaches directed against WT spike should be effective against the D614G strains. In addition, these data support the hypothesis that early in the pandemic S-trimer reorganization favors transmission over-sensitivity to neutralization, a phenotype that might be expected to emerge as a new virus spreads through a large naive population and then undergoes new evolutionary change as herd immunity increase with time.

In summary, our data support the critical need to periodically review SARS-CoV-2 contemporary isolates across the globe and identify the emergence of new variants with increased transmission and pathogenesis and/or altered antigenicity, especially as levels of human herd immunity and interventions alter the selective forces that operate on the genome. Our data suggest that vaccines encoding the ancestral D614 S glycoprotein will elicit robust neutralization titers against contemporary G614 isolates, supporting continued development of existing vaccine formulations.

## Acknowledgements

This work was supported by grants from the U.S. National Institutes of Health, including R01-AI110700, U54-CA260543, U01-AI151797, and R01-AI069274 along with contracts HHSN272201700036I and HHSN272201400008C. This work was supported in part by the Japan Agency for Medical Research and Development under grant numbers JP19fk0108113, JP19fm0108006, JP20fk0108104 and JP19fk0108110. This project was also supported by the North Carolina Policy Collaboratory at the University of North Carolina at Chapel Hill with funding from the North Carolina Coronavirus Relief Fund established and appropriated by the North Carolina General Assembly. We thank Ms. Yuko Sato at the National Institute of Infectious Diseases for technical support with pathological analysis. We are grateful to Adimab LLC. for providing us SARS-CoV-2 nAbs S309, REGN10933, REGN10987 and JS016. We thank Kenichi Okuda and Takafumi Kato at UNC Marsico Lung Institute for technical support on primary cells.

## Material and Method

### Antibodies

Monoclonal SARS-CoV-2 RBD-binding neutralizing antibodies (nAb) B38 and H4 were synthesized at UNC Protein Expression and Purification core based on previously reported protein sequences (*27*). The nAbs S309, REGN10933, REGN10987 and JS016 were reported previously (*28, 29*) and were kindly provided by Adimab LLC. Serum samples collected from BALB/c mice vaccinated with WA1 spike protein (D-form) were generated in our laboratory previously (*4, 14*). Monoclonal antibody targeting the cytoplasmic tail of SARS-CoV-2 S protein was purchased from Abcam (ab272504). Polyclonal antibodies targeting the SARS-CoV N protein PA1-41098 and ANT-180 were purchased from Invitrogen and Prospec, respectively. Mouse antiserum targeting SARS-CoV-2 nucleocapsid protein was produced in our laboratory as described previously (*14*).

### Cells and viruses

Simian kidney cell lines Vero-81 (ATCC # CCL81), Vero-E6 (ATCC # CRL1586) were maintained in Eagle’s Minimum Essential Medium (Gibco) supplemented with 10% fetal calf serum (FBS, Hyclone). Huh7 and A549-ACE2 cells were maintained in Dulbecco’s Modified Eagle Medium (Gibco) with 10% FBS. A clonal A549-ACE2 stable cell line was generated by overexpressing human ACE2 in the A549 cell line (ATCC # CCL185) using the Sleeping Beauty Transposon system. Generation of primary human pulmonary cell cultures was described previously (4). Primary human nasal epithelial cells (HNE) were collected from healthy volunteers by curettage under UNC Biomedical IRB-approved protocols (#11-1363 and #98-1015). Human bronchial epithelial [large airway epithelial (LAE)] and bronchiolar [small airway epithelial (SAE)] cells were isolated from freshly excised normal human lungs obtained from transplant donors with lungs unsuitable for transplant under IRB-approved protocol (#03-1396) and cultured in air liquid interface (ALI) media, as previously described (*4, 30*). SARS-CoV-2 WA1 molecular clone, WT and nLuc viruses were generated previously (*4, 14*). To generate the D614G and D614G-nLuc variants, the amino acid substation was introduced into the S gene in the plasmid F and coupled with plasmid G with or without nLuc insertion in the ORF7a. Then, the seven genomic cDNA fragments spanning the entire SARS-CoV-2 genome were digested, purified and ligated. Full-length RNA was transcribed and electroporated into Vero E6 cells. Virus stocks were verified by Sanger sequencing. All viral infections were performed under biosafety level 3 (BSL-3) conditions at negative pressure, and Tyvek suits connected with personal powered-air purifying respirators.

### nLuc virus entry assay

Monolayers of Vero-E6, Vero-81, A549-ACE2 and Huh7 cells were cultured in black-walled 96-well plates (Corning 3904) overnight. The cells were infected with WT-nLuc or D614G-nLuc viruses at MOI of 0.1. After incubation for 1h, inocula were removed, and the cells were washed two times with PBS and maintained in DMEM containing 5% FBS and the mixture of SARS-CoV-2 nAbs REGN10933, REGN10987 and JS016 at concentration of 1000 times of IC_50_ for each. After incubation at 37°C for 48h, viral infection was quantified using nLuc activity via Nano-Glo Luciferase Assay System (Promega) according to the manufacturer specifications.

### SARS-CoV-2 neutralization assay

Vero E6 cells were plated at 20,000 cells per well in black-walled 96-well plates (Corning 3904). Mouse serum samples were tested at a starting dilution of 1:20 and mAb samples were tested at a starting dilution 30 to 0.1 μg/ml and were serially diluted 3-fold up to eight dilution spots. Diluted antibodies or sera were mixed with 189 PFU/well WT-nLuc or D614G-nLuc virus, and the mixtures were incubated at 37°C for 1 hour. Following incubation, growth media was removed, and virus-antibody mixtures were added to the cells in duplicate. Virus-only controls were included in each plate. Following infection, plates were incubated at 37°C with 5% CO2 for 48h. After the 48h incubation, cells were lysed, and luciferase activity was measured via Nano-Glo Luciferase Assay System (Promega) according to the manufacturer specifications. Neutralization titers were defined as the sample dilution at which a 50% reduction in relatively light unit (RLU) was observed relative to the average of the virus control wells.

### WT and D614G competition assay and BstCI digestion

LAE cultures from one donor were infected with MOI of 0.5 of WT and D614G mixture at 1:1 and 10:1 ratios. Following 1h incubation, the cultures were washed three times with PBS and cultures for 72h in the air-liquid interface (ALI) condition. To passage the progeny viruses, 100uL PBS was added to each LAE surface for 10 min incubation and was added to naïve cultures surface for infection. The virus samples were continuously passaged three times in LAE culture, and cellular RNA samples from the 3^rd^ passage were extracted using TRIzol reagent (Thermo Fisher). A 1547bp fragment containing the D614G site was amplified from each RNA samples by RT-PCR using primer set: 5’-GTAATTAGAGGTGATGAAGTCAGAC-3’ and 5’-GAACATTCTGTGTAACTCCAATACC-3’. The amplicon was purified by agarose gel electrophoresis and digested with BstCI restriction enzyme (NEB) overnight. The digested products were analyzed on agarose gel electrophoresis.

### hACE2 mice infection and titration

Mouse study was performed in accordance with Animal Care and Use Committee guidelines of the University of North Carolina at Chapel Hill. *HFH4-hACE2* transgenic mice were bred and maintained at UNC. Mice were infected with 10^5^ PFU of WT or D614G viruses intranasally under ketamine/xylazine anesthesia. At indicated timepoints, a subset of mice were euthanized by isoflurane overdose, and tissue samples were harvested for viral titer analysis. The right caudal lung lobe, brain and nasal turbinates were taken for titer and stored at −80 °C until homogenized in 1mL PBS and titrated by plaque assay. Briefly, the supernatants of homogenized tissue were serially diluted in PBS, 200 μL of diluted samples were added to monolayers of Vero-E6 cells, followed by agarose overlay. Plaques were visualized by day 2 post staining with neutral red dye.

### Whole-mount immunostaining and imaging

WT or D614G-infected LAE ALI cultures were fixed twice for 20 minutes in 4% formaldehyde in PBS and stored in PBS. The SARS-CoV-2 N antigen was stained with polyclonal rabbit anti-SARS-CoV N protein (Invitrogen PA1-41098, 0.5 ug/mL),and using species-specific secondary antibodies as previously described (*4*). The cultures were also imaged for α-tubulin (Millipore MAB1864; 3ug/mL) and MUC5AC (Thermo Scientific 45M1; 4ug/mL) as indicated. Filamentous actin was localized with phalloidin (Invitrogen A22287), and nuclei was visualized with Hoechst 33342 staining (Invitrogen). An Olympus FV3000RS confocal microscope in Galvo scan mode was used to acquire 5-channel Z stacks by 2-phase sequential scan. Representative stacks were acquired and are shown as Z-projections and XZ cross sections to distinguish individual cell features and to characterize the infected cell types. ImageJ was used to measure the relative apical culture surface covered by multiciliated cells.

### Western blot analysis of spike protein cleavage

Exocellular SARS-CoV-2 virions were collected from WT or D614G infected LAE culture by gently washing intact apical surface with 100uL PBS. Samples from the triplicated cultures were pooled, lysed with modified RIPA buffer and inactivated at 98 °C. Protein samples were electrophoresed in 4-20% continuous SDS-PAGE gel (Bio-Rad) and transferred on to a PVDF membrane (Bio-Rad). SARS-CoV-2 S protein was probed using a mAb targeting SARS-CoV-2 S mAb (Abcam, ab272504) and the N protein was probed using a mouse antiserum produced in our laboratory. The N protein was used as a loading control for the Western blot.

### EM imaging and Spike quantification

WT or D614G infected primary cell cultures were submerged in fixative (4% paraformaldehyde, 2.5% glutaraldehyde and 0.1 M sodium cacodylate) overnight. For SEM, samples were rinsed, fixed with 1% OsO4 (Electron Microscopy Sciences) in perfluorocarbone FC-72 (Thermo Fischer) solution for 1 hour. After dehydration and mounted on aluminum planchets, samples were imaged using a Supra 25 field emission scanning electron microscope (Carl Zeiss Microscopy). For TEM, fixed samples were rinsed and post-fixed with potassium-ferrocyanide reduced osmium (1% osmium tetroxide/1.25% potassium ferrocyanide/0.1 sodium cacodylate buffer. The cells were dehydrated and embedment in Polybed 812 epoxy resin (Polysciences). The cells were sectioned perpendicular to the substrate at 70nm using a diamond knife and Leica UCT ultramicrotome. Ultrathin sections were collected on 200 mesh copper grids and stained with 4% aqueous uranyl acetate followed by Reynolds’ lead citrate. Samples were observed using a JEM-1230 transmission electron microscope operating at 80kV and images were taken using a Gatan Orius SC1000 CCD camera (Gatan). The number of spikes on each virion projection was quantified using ImageJ software. SEM images of infected cultures from 3 donors were imaged, at least 10 different micrographs (>100k X) were analyzed using the multi-point counting tool on individual virions.

### Hamster infection, tissue collection, and transmission studies

Hamster studies were performed in accordance with Animal Care and Use Committee guidelines of the University of Wisconsin-Madison. Syrian hamsters (females, 4-6 weeks old) were purchased from Envigo (Madison, WI) and allowed to acclimate for a minimal of three days at BSL-3 agriculture containment at the Influenza Research Institute (University of Wisconsin). Hamsters were infected with 10^3^ PFU of WT or D614G viruses intranasally under isoflurane anesthesia. At the indicated timepoints, a subset of hamsters were euthanized by deep anesthesia by isoflurane inhalation and cervical dislocation and tissues samples were harvested for virus titer and histopathology analysis. Weights of one group of hamsters were recorded for 14 days after infection.

To evaluate indirect virus transmission between hamsters, groups of hamsters (n=8 per group) were infected with 10^3^ PFU of WT or D614G viruses intranasally under isoflurane anesthesia. Infected animals were placed in specially designed cages (Figure S2B) inside an isolator unit (*16*). Twenty-four hours later, naïve hamsters were placed in the other side of the cage with 5 cm separation by a double-layered divider to allow free air flow. The isolator unit provided one directional airflow; therefore, the infected hamsters were placed in the front of the isolator unit. Metal shroud were placed over the cages so only the front and back of the cage was open. Nasal washes were collected at 3-day intervals for the infected hamsters and 2-day intervals for the exposed animals starting on day 2 after infection or exposure (Figure S2A).

### Pathological examination

Tissues fixed for at least seven days in 10% formalin were trimmed and embedded in paraffin. The paraffin blocks were cut into 3 μm-thick sections and mounted on silane-coated glass slides. One section from each tissue sample was stained using a standard hematoxylin and eosin procedure. To detect SARS-CoV-2 Nucleocapsid protein in immunohistochemistry (IHC), tissue sections were incubated with a rabbit polyclonal antibody (Prospec, ANT-180) as the primary antibodies, and peroxidase-labeled polymer-conjugated anti-rabbit immunoglobulin (EnVision/HRP, DAKO) as the secondary antibody. Immunostaining was visualized by 3,3’-diaminobenzidine tetrahydrochloride staining. Hematoxylin (Modified Mayer’s) was used as a nuclear counterstain for IHC.

---

**Figure S1.**
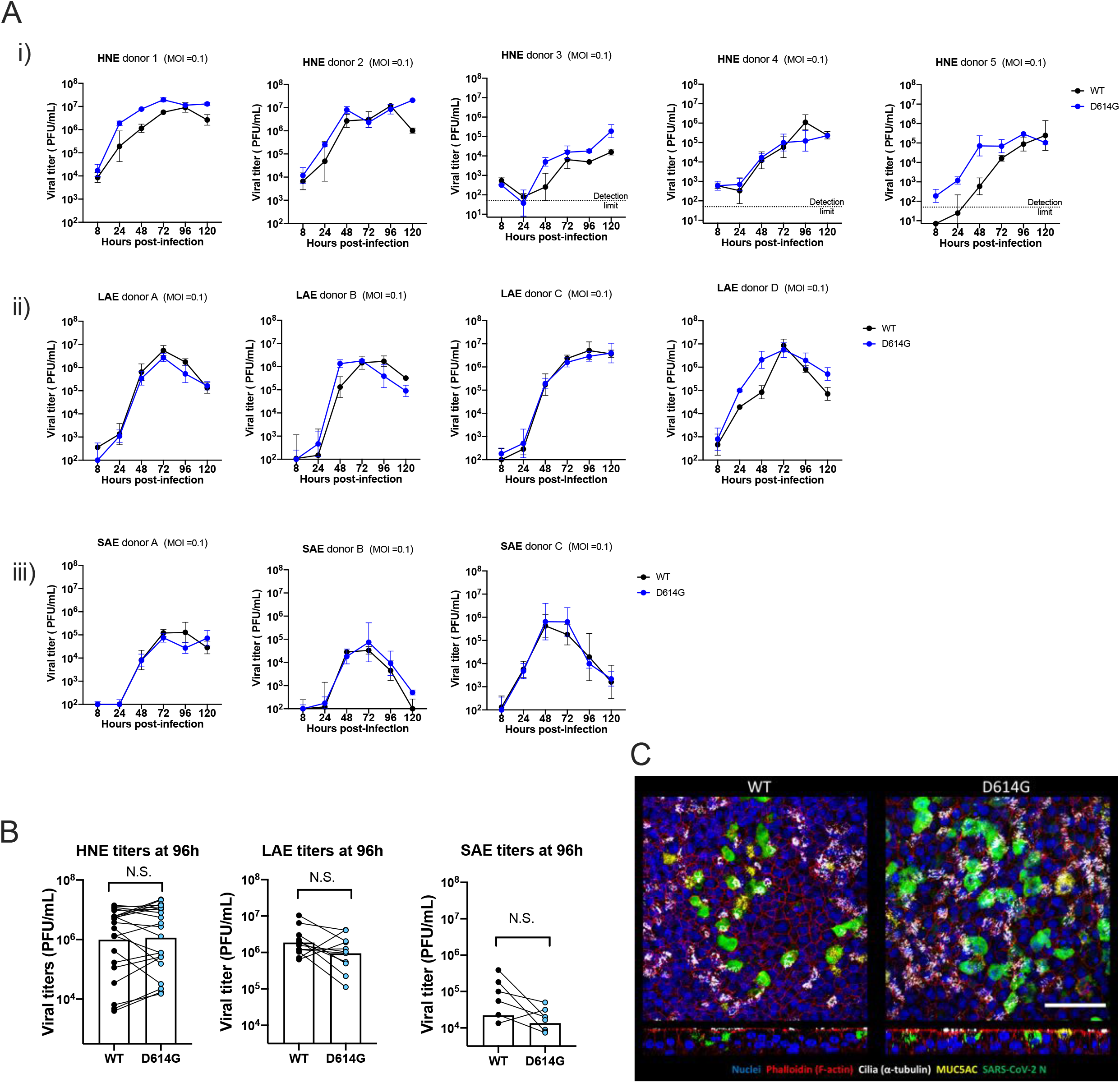
Additional data of WT and D614G infected primary human airway epithelial cells. A. Growth curves of the two viruses in individual primary nasal (i), large airway (ii) and small airway (iii) epithelial cells relating to Fig 1 D to F, MOI = 0.1; plaque assay detection limit: 1.7 log_10_PFU/mL. B. comparison of WT and D614G titer at 96h on HNE, LAE and SAE. C. Whole-mount staining of WT and D614G infected LAE cultures, blue: Hoechst (nuclei), red: phalloidin (F-actin), white: cilia (α-tubulin); yellow: MUC5AC, Green: SARS-CoV-2 N protein, scale bar: 50μm.

**Figure S2.**
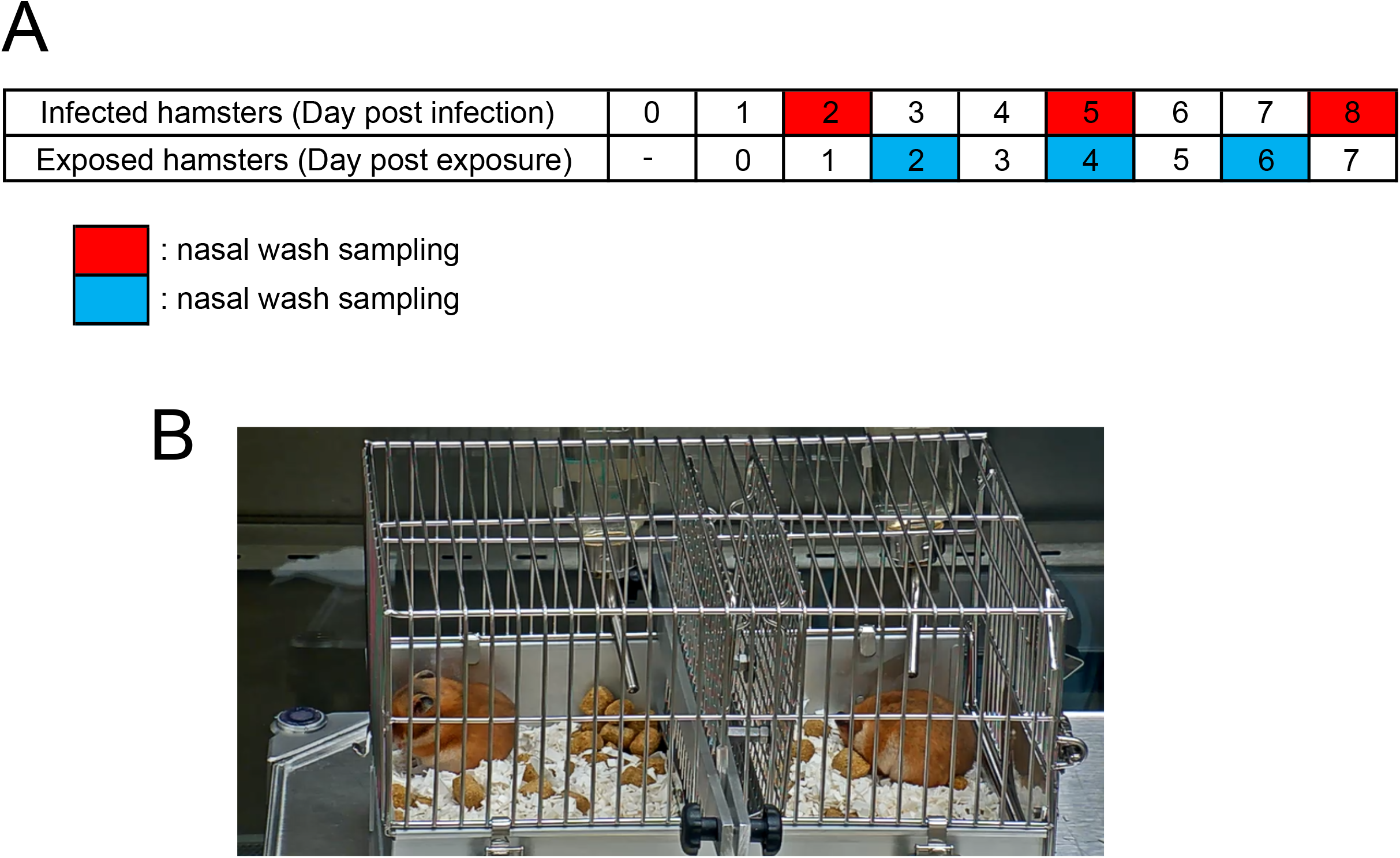
Experimental design of hamster transmission study. A, Timeline of nasal wash sampling from infected and exposed animals. B. Image of hamsters in the cage in transmission study.

